# Characterising Genetic Diversity in Cassava Brown Streak Virus

**DOI:** 10.1101/455303

**Authors:** Stephen M Crotty, Adam B Rohrlach, Joseph Ndunguru, Laura M Boykin

## Abstract

Plant viruses represent a significant threat to food security for many global populations. Cassava Brown Streak Virus (CBSV) causes immense damage to cassava crops in Eastern, Central and Southern Africa. The eradication of CBSV is a difficult challenge, as it has been shown to be fast-evolving and it is transmitted by flying insects that are ubiquitous in cassava growing regions. In this paper we demonstrate the ability of two new developments in bioinformatics that can be used to increase our understanding of CBSV and ultimately inform strategies for its combat. We reconstruct the phylogeny of 29 whole-genome virus isolates using the GHOST model. This phylogeny identifies three distinct clades among the viruses and highlights a section of the genomes that is highly influential in their divergence. We also perform Multiple Correspondence Analysis on the alignment which is consistent in recovering the three clades, and offers insight on the significance of the influence of a variety of external variables on the evolution of the viruses. Knowledge and information from this analysis will be used as a base on which to formulate sustainable Cassava Brown Streak Disease (CBSD) management strategies in Africa.

## INTRODUCTION

It is difficult to overemphasize the importance of agriculture to people living in sub-Saharan Africa. The population of sub-Saharan Africa exceeds one billion people, and more than 60% of the population live in rural areas (United Nations, 2015). Beyond the obvious benefit of nutrition, agriculture is also the primary source of income for the majority of sub-Saharan African families, and so access to essentials such as health care and education depend indirectly on agriculture. As such, the productivity of the agricultural industry in a given country and year is a key indicator of the health and wellbeing of the population and the economy.

One of the major threats to agricultural productivity comes in the form of crop pests and diseases. Of particular significance are the viruses of the genus *Ipomovirus.* They have the potential to devastate crops, and they infect some of the most economically important, commonly grown food staples across sub-Saharan Africa, such as cassava and sweet potato. Cassava Brown Streak Disease (CBSD) is caused by two closely related *Ipomovirus* species, Cassava Brown Streak Virus (CBSV) and Ugandan Cassava Brown Streak Virus (UCBSV). CBSD is widespread and causes significant reduction in both the quality and the yield of cassava crops, making it a strong barrier to food security and economic prosperity in the regions in which it is grown.

Efforts to combat CBSV and UCBSV are ongoing, but there are many challenges. The viruses are transmitted between plants courtesy of whiteflies *(Bemesia tabaci)* and other potential vectors (Ateka et al., 2017), so it is critical that infected plants are removed from crops swiftly. This indicates a critical need for early detection and positive diagnosis. However, traditional sequencing methods such as PCR and Sanger sequencing are slow and costly, with a positive diagnosis taking weeks or months, far too late to prevent the virus from spreading. A recent advance in sequencing technology has seen researchers use nanopore sequencing to take samples from plants, and then sequence and identify pathogens in just a few hours, without the need to leave a farmer’s field (Boykin et al., 2018). This enables the swift quarantine of infected material, protecting the rest of their crop and neighbouring crops.

Efforts to isolate and destroy infected plant material are on the frontline of the battle against CBSV and UCBSV, but they are reactive strategies in that they combat the virus after plant infection has already occurred. It is necessary to simultaneously pursue proactive strategies, such as breeding resistant forms of cassava. This is not straightforward though, as it has already been shown that these viruses are fast-evolving (CBSV moreso than UCBSV) (Alicai et al., 2016). Mbewe et al. (2017) showed that in the gene tree of the P1 gene, a third distinct clade is found in addition to CBSV and UCBSV, which they tentatively labelled Tanzanian CBSV (CBSV-TZ). Critical to winning the fight against CBSD is ongoing research into the evolutionary forces acting on these viruses, at both the molecular level and geographically.

We made use of two new bioinformatics tools to analyse a sequence alignment consisting of 29 whole virus genomes, 14 CBSV and 15 UCBSV. We first performed phylogenetic inference under the GHOST model of sequence evolution (Crotty et al., 2017). The GHOST model is a mixture model which enables the fitting of multiple classes to a single alignment. The tree topology is common across all classes, but each class has its own set of branch lengths and model parameters. This enables subtle phylogenetic signals to be extracted from the data, which is especially useful when there exists a clear dominant signal, such as that distinguishing the two virus species in our alignment. We also applied Multiple Correspondence Analysis (MCA) to the alignment, following the method outlined in Rohrlach et al. (2018). They demonstrated MCA as an effective tool for decomposing the variability within categorical sequence data for visualization in two dimensions. Furthermore, they showed that it was possible to meaningfully associate the genetic diversity with geographical coordinates, allowing for informed hypotheses relating to the path of evolution to be constructed.

## METHODS

### Data

The alignment consisted of 14 CBSV and 15 UCBSV whole genome sequences. The alignment was obtained from Alicai et al. (2016), in which details of its assembly can be found.

### Phylogenetic Analysis

We used IQ-TREE (Nguyen et al., 2015) to fit a GHOST model to the sequence alignment. We used the model selection procedure outlined in Crotty et al. (2017) to choose the model of sequence evolution and number of classes. After performing the inference we analysed the site-wise probabilities of evolving under each inferred class to identify influential sites, contiguous regions and genes within the genome.

### Multiple Correspondence Analysis

We conducted MCA following the procedure outlined in Rohrlach et al. (2018). Due to the fast rate of evolution present we removed all singleton sites: sites where the nucleotide was conserved across all but one taxon. We carried out MCA on the entire sequence alignment, as well as on the 14 CBSV taxa and 15 UCBSV taxa separately.

## RESULTS

### Phylogenetic Analysis

#### Model selection

In order to determine the optimal model of sequence evolution and number of classes when fitting the GHOST model, we experimented with a wide variety of substitution models, fitting each one with between two and twelve classes. We then identified the best fitting combinations according to both Bayesian Information Criterion (BIC) (Schwarz et al., 1978) and Akaike’s Information Criterion (AIC) (Akaike, 1974). Results can be found in Supplementary Figures S1 and S2. The smallest BIC score was obtained using the General Time Reversible (GTR) model and three classes, while the smallest AIC score was obtained using GTR with ten classes. It is not unexpected to see such a discrepancy with the parameter-rich GHOST model. BIC places a relatively heavy penalty on additional parameters and is therefore prone to underfitting, whereas AIC imposes a relatively light penalty and is therefore prone to overfitting (Burnham and Anderson, 2003; Posada and Buckley, 2004; Dziak et al., 2012).

Examining the recovered trees from the 3-class GHOST model (Figure 1) preferred by BIC, distinct and reasonable biological interpretations can be made for all three classes. The first class has very short branch lengths, indicating strong conservation of nucleotides across all taxa. Since these viruses are closely related, we can think of this class as capturing the component of the phylogenetic signal that is common to CBSV and UCBSV. The second class shows only one branch length of any significance, that separating the CBSV and UCBSV clades. Within the two clades the branch lengths are again very short, indicating that this class captures the component of the phylogenetic signal relating to the divergence of CBSV and UCBSV. The third class distinguishes itself from the second by a clade of four CBSV replicates that diverges from both UCBSV and the remaining CBSV replicates.

**Figure 1:**
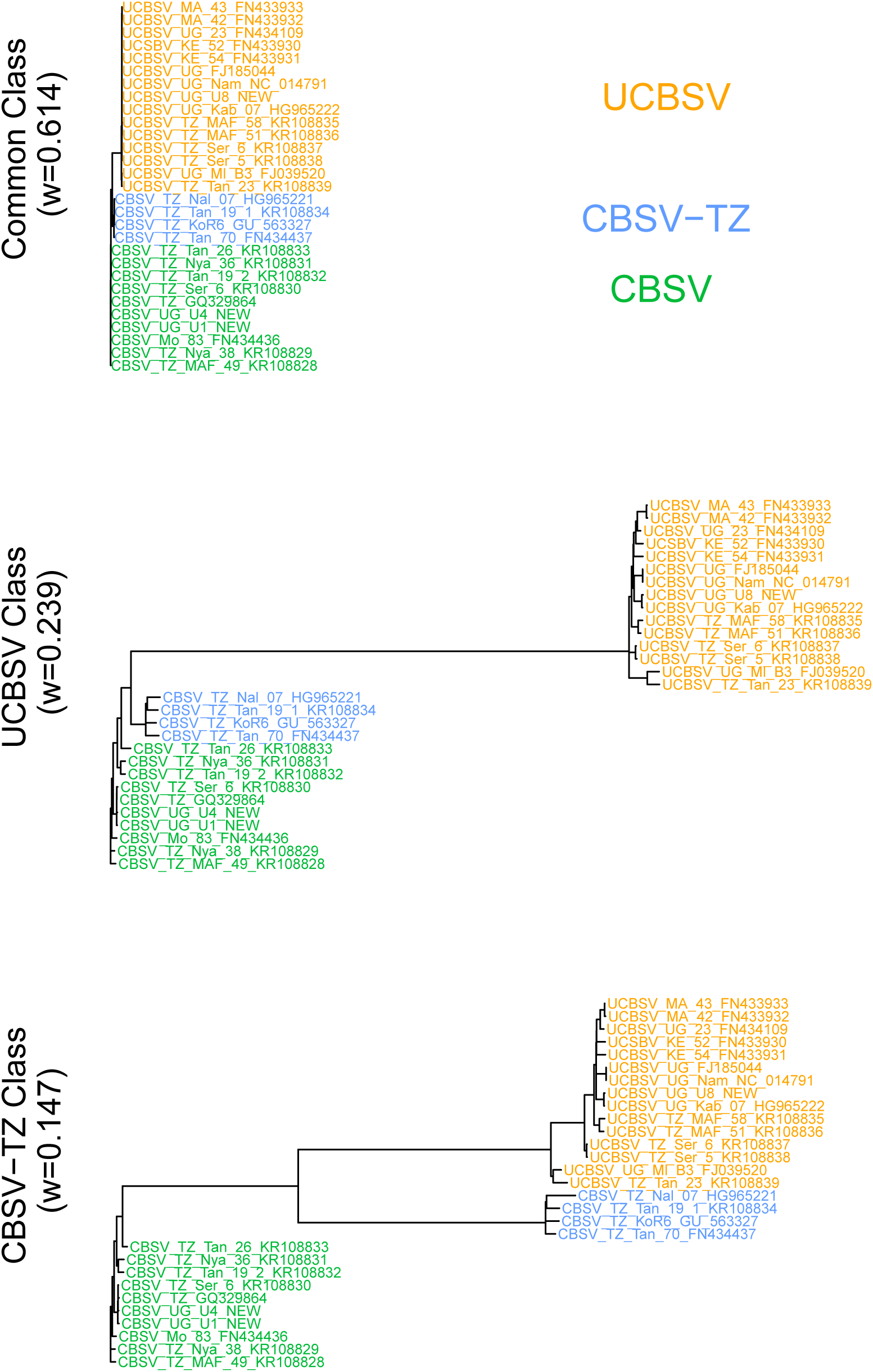
Trees inferred by IQ-TREE using the GHOST model with three GTR classes.

Unlike the trees of the 3-class model, the trees of the 10-class model (Supplementary Figure S3) preferred by AIC bear the hallmarks of overfitting, as described in Crotty et al. (2017). Many of the classes bear strong similarity to each other, most noticeably the first three classes all appear strongly conserved across all taxa, much like the first class of the 3-class model. In fact, it is not unreasonable to suggest that all 10 class trees could be loosely categorised as falling into one of the three categories defined above by the trees of the 3-class model. No significant new phylogenetic signal appears to be captured by this model, beyond those which were found by the 3-class model. Consequently, we concluded that the 3-class GHOST model with a GTR model of sequence evolution provided the best fit to this alignment.

#### Identifying regions of interest

Having chosen the optimal GHOST model, we can use the results to identify regions within the genome that may be influential in the patterns of evolution that we observe. For convenience, we refer to the first class as the Common Class, since there is no significant divergence among any of the taxa; we refer to the second class as the UCBSV Class, since this class is characterised by the divergence between the CBSV and UCBSV replicates; and we refer to the third class as the CBSV-TZ Class. The reason for this is that the four CBSV replicates that diverge from the remainder in this class are also present in the study of the P1 gene (Mbewe et al., 2017) referred to earlier. These four replicates are found in the third clade which they tentatively labelled CBSV-TZ, and so it seems probable that the phylogenetic signal captured in the third class is related to the divergence of CBSV-TZ from CBSV. It is worth noting here that we do not make the claim that CBSV-TZ is a new species of CBSV, but use these terms for convenience in labelling the classes, and consistency with the work of Mbewe et al. (2017). Further research is required in order to establish CBSV-TZ as a new vspecies of the virus.

The contribution of a particular site in the alignment to the likelihood function of a mixture model such as GHOST is simply the weighted sum of the partial likelihoods of that site under each class. We can make use of these partial likelihoods to determine for every site the probability distribution of evolving under each of the classes. Put simply, we can look at each site and identify the probability with which it belongs to any of the classes. This allows us to locate particular sites, contiguous regions and genes that belong to the CBSV-TZ class with high probability, thereby being identified as influential in the evolutionary divergence of CBSV-TZ and CBSV. In Table 1 we list the ten genes that consititute the genome of the viruses, the length of the genes in nucleotides as well as in percentage of the genome, and finally the percentage contribution of the genes towards the CBSV-TZ Class. Most apparent from the Table 1 is that the P1 gene appears to be of strong influence in the divergence of CBSV-TZ and CBSV. The 1086 sites that make up the P1 gene, representing 12.41% of the entire genome, account for more than 22% of the weight of the CBSV-TZ Class. The remaining genes contribute to the CBSV-TZ Class approximately in proportion with their relative length in the genome, with the exception of the CP gene which constitutes 12.96% of the genome but only accounts for 6.57% of the CBSV-TZ Class.

**Table 1:** A list of the 10 genes that make up CBSV and UCBSV. Also shown is the size of each gene in units of nucleotides and as a percentage of the entire genome. The final column shows the percentage of the CBSV-TZ Class attributable to each gene.

| Gene | Size | % Genome | % CBSV-TZ Class |
| --- | --- | --- | --- |
| P1 | 1086 | 12.41 | 22.08 |
| P3 | 882 | 10.08 | 10.37 |
| 6K1 | 156 | 1.78 | 1.91 |
| CI | 1890 | 21.61 | 20.27 |
| 6K2 | 156 | 1.78 | 1.86 |
| Vpg | 558 | 6.38 | 7.19 |
| NIa | 702 | 8.03 | 8.45 |
| NIb | 1506 | 17.22 | 15.61 |
| Ham1 | 678 | 7.75 | 5.69 |
| CP | 1134 | 12.96 | 6.57 |
| Total | 8748 | 100 | 100 |

In Figure 2, we show the cumulative probability of sites in the P1 gene belonging to the CBSV-TZ Class. It displays graphically the observation made from Table 1, that the average gradient of the P1 gene (represented by the red dotted line) is nearly double that of the average gradient across the entire genome (represented by the blue dotted line). Additionally, Figure 2 also demonstrates that the contribution to the CBSV-TZ Class is not even across the entire gene. Compared to the average for the entire gene, there is a noticeable increase in gradient, roughly between nucleotide 70 and 170. We therefore surmise this to be the region of highest influence within the gene of highest influence. Figures 3-5 show the alignment for the first 80 amino acids of the P1 gene. Examining the region of interest identified above (amino acids 23 to 57, approximately), we notice in Figure 3 that there is strong conservation among the CBSV isolates, yet the UCBSV and CBSV-TZ isolates are relatively diverged. Similarly, Figure 4 shows that there is also strong conservation among the 15 UCBSV isolates in this region. In contrast, Figure 5 shows comparatively little conservation among the four CBSV-TZ isolates. In summary, while all three strains are strongly divergent from each other in this region, within group genetic variability is low in CBSV and UCBSV but high in CBSV-TZ.

**Figure 2:**
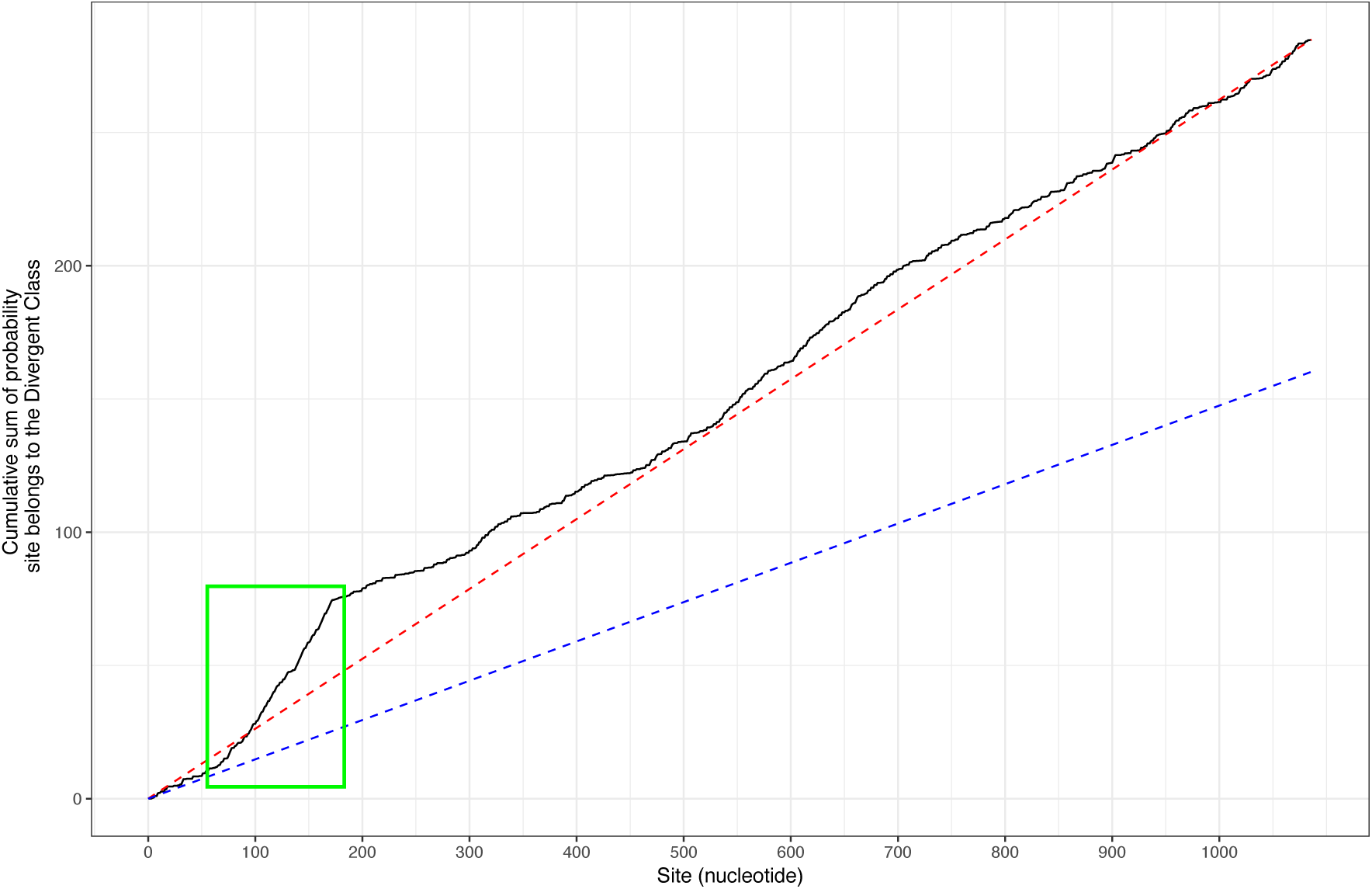
A cumulative sum plot of the probability of sites belonging to the CBSV-TZ Class, for the P1 gene. The average gradient in any contiguous region is representative of the contribution of this region to the CBSV-TZ Class. The red dashed line indicates the average probability of the sites in the P1 gene belonging to the CBSV-TZ Class. The blue dashed line indicates the average probability of all sites in the genome belonging to the CBSV-TZ Class. The green box highlights a section of the gene in which the gradient is particularly steep, indicating that this region contributes strongly to the CBSV-TZ Class.

**Figure 3:**
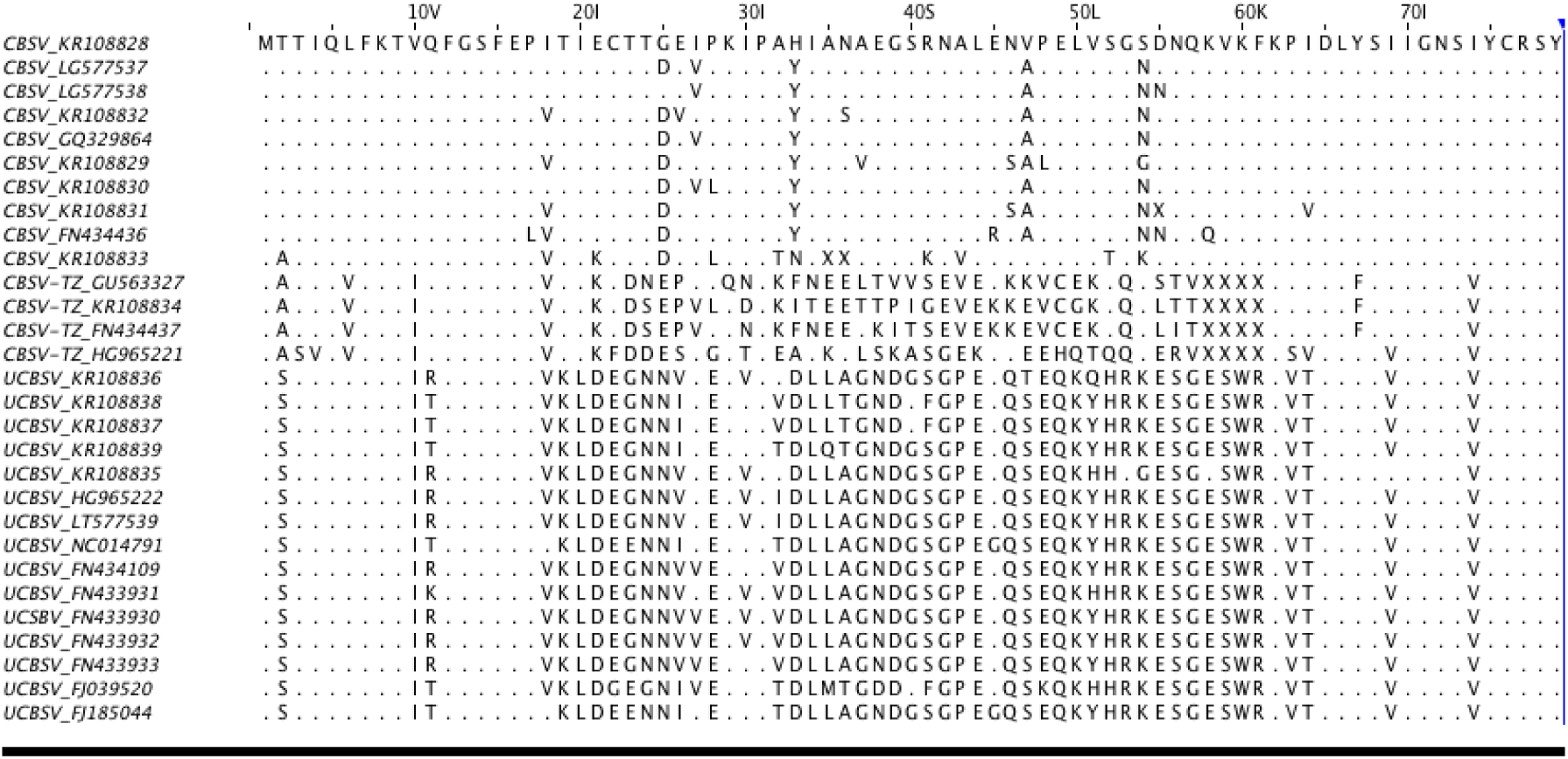
The first 80 amino acid residues in the alignment of the P1 gene of the 29 CBSV and UCBSV sequences. The top CBSV sequence is used as a reference, with the conserved residues represented as dots.

**Figure 4:**
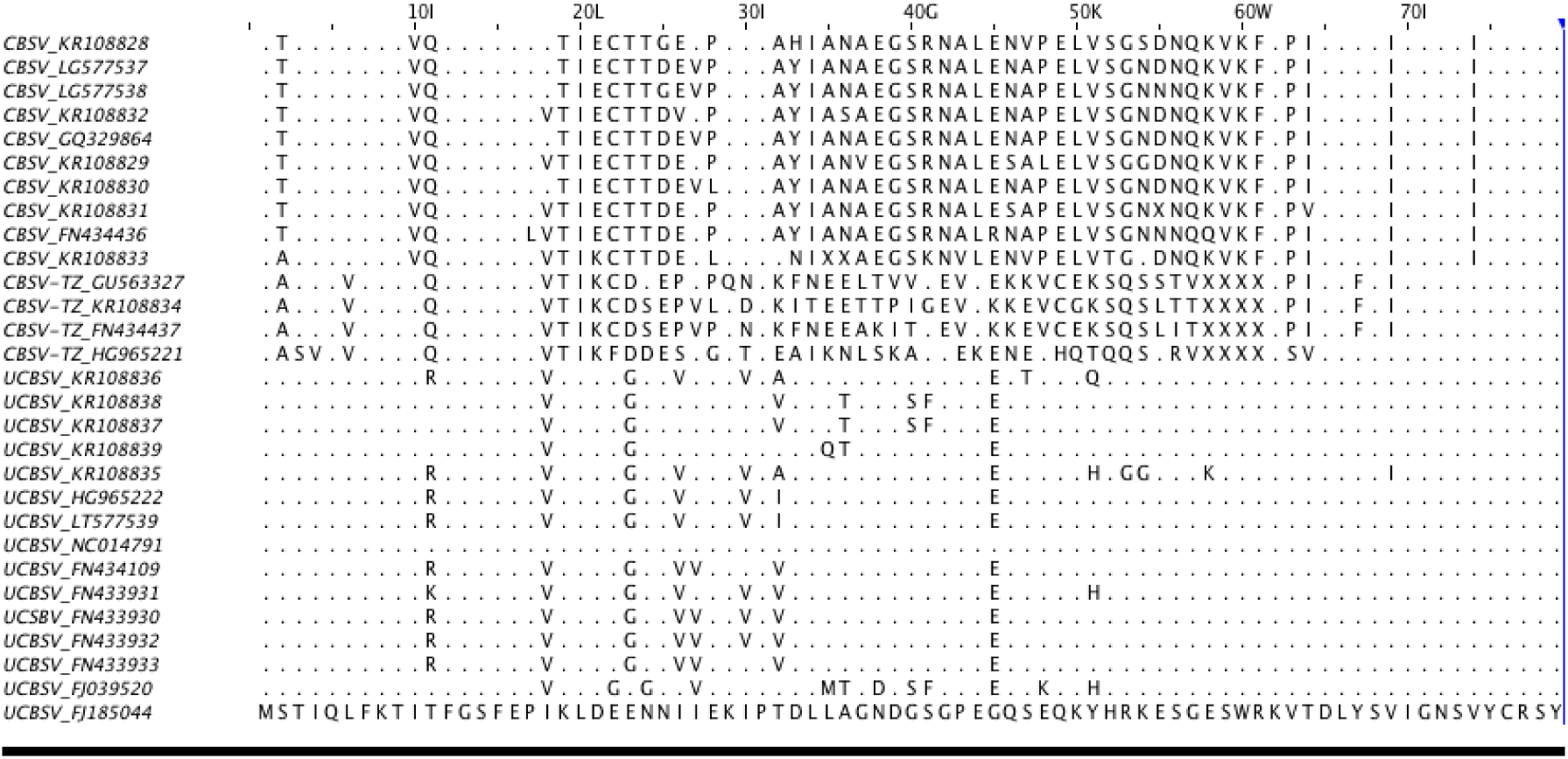
The first 80 amino acid residues in the alignment of the P1 gene of the 29 CBSV and UCBSV sequences. The bottom UCBSV sequence is used as a reference, with the conserved residues represented as dots.

**Figure 5:**
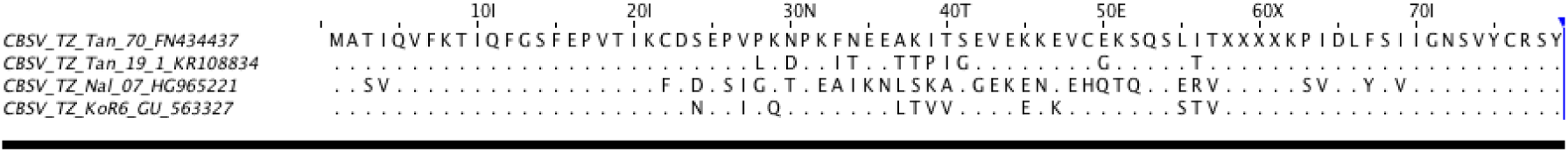
The first 80 amino acid residues in the alignment of the P1 gene of the 4 CBSV-TZ sequences. The top CBSV-TZ sequence is used as a reference, with the conserved residues represented as dots.

#### MCA analysis

When applied to the entire dataset, Figure 6 shows that the distinction between the three strains is the dominant feature that is captured by MCA. The first principal dimension clearly distinguishes between CBSV (including the four CBSV-TZ sequences), while the second dimension distinguishes between CBSV and CBSV-TZ. The signals that distinguish the three strains are so dominant that it is difficult to observe anything additional from performing MCA on the entire dataset. To search for more nuanced patterns in the alignment, we performed MCA on meaningful subsets of the genomes.

**Figure 6:**
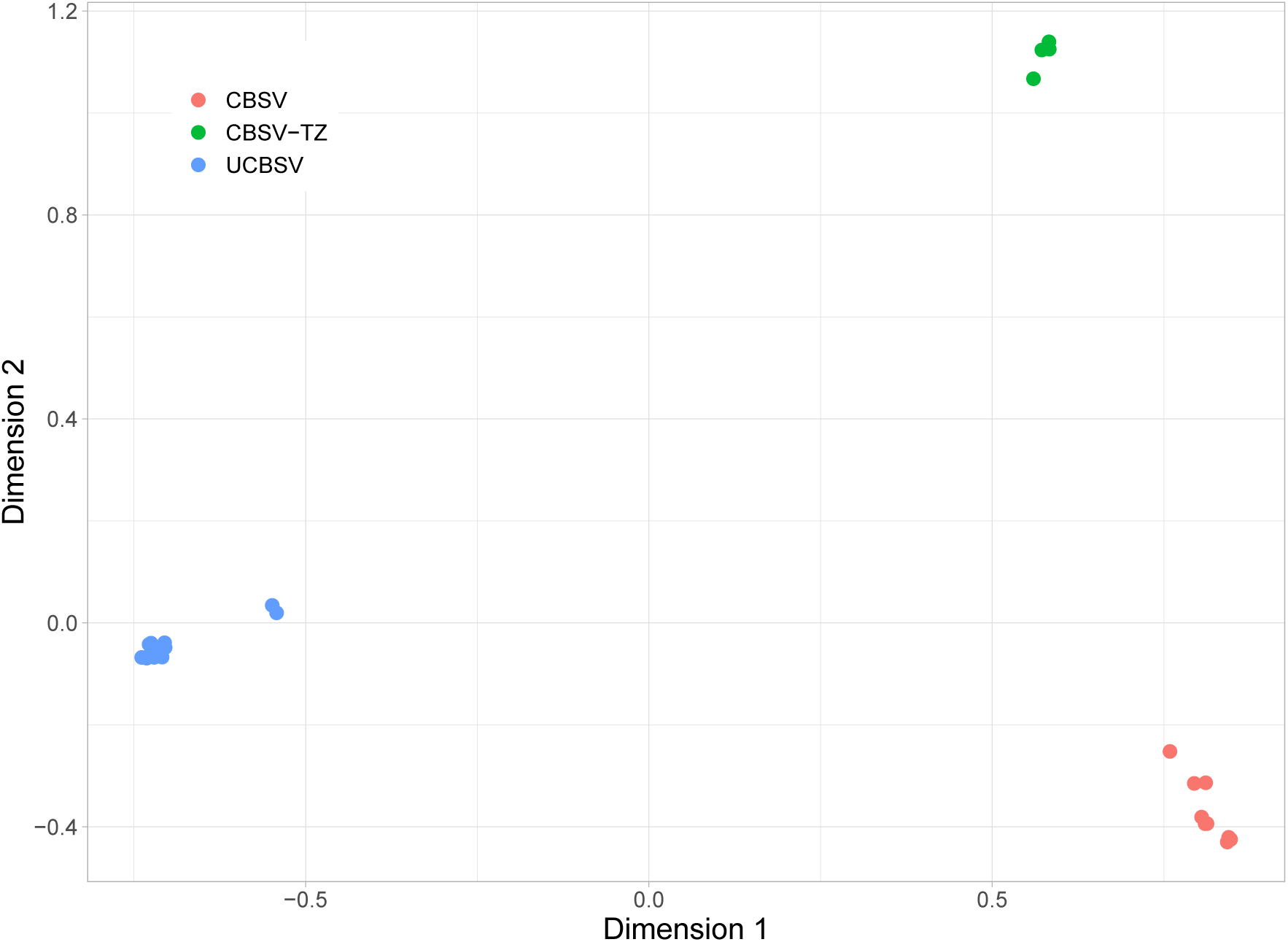
Results of Multiple Correspondence Analysis (MCA) on the full alignment. The three distinct clusters correspond to the three strains of CBSV.

Figure 7 displays four scatterplots of the first two dimensions of a MCA performed on the 14 CBSV isolates. Each scatterplot is coloured according to a different variable, to aid in highlighting any patterns in the data. Once again, the first principal dimension captures the most dominant feature, the distinction between the CBSV and CBSV-TZ strains. A clear pattern is not obvious in Figures 7 a) and b), which colour the points according to the geographic origin of the isolates (latitude and longtitude). The points in Figure 7 c) are coloured according to the sample collection date. Most samples in the dataset were collected either in 2013 (15 of 29), or in 2006 - 2008 (11 of 29). We therefore used binary colouring according to these two groups. The striking feature of Figure 7c), is that most of the CBSV-TZ isolates were collected prior to 2009 (3 of 4), whereas most of the CBSV isolates (8 of 10) were collected in 2013. That is, 60% of the samples collected before 2009 were CBSV-TZ, whereas only 11% of the samples collected in 2013 were CBSV-TZ. A similar pattern emerges when observing Figure 7 d), in which points are classified as either lowlands (less than 1000m above sea level) or highlands (more than 1000m above sea level). The threshold was chosen as historically, CBSD was not found in areas that were greater than 1000m above sea level (Alicai et al., 2007). We see from Figure 7 d) that all of the CBSV-TZ isolates came from lowland areas, while the CBSV isolates are approximately evenly split between lowland and highland areas.

**Figure 7:**
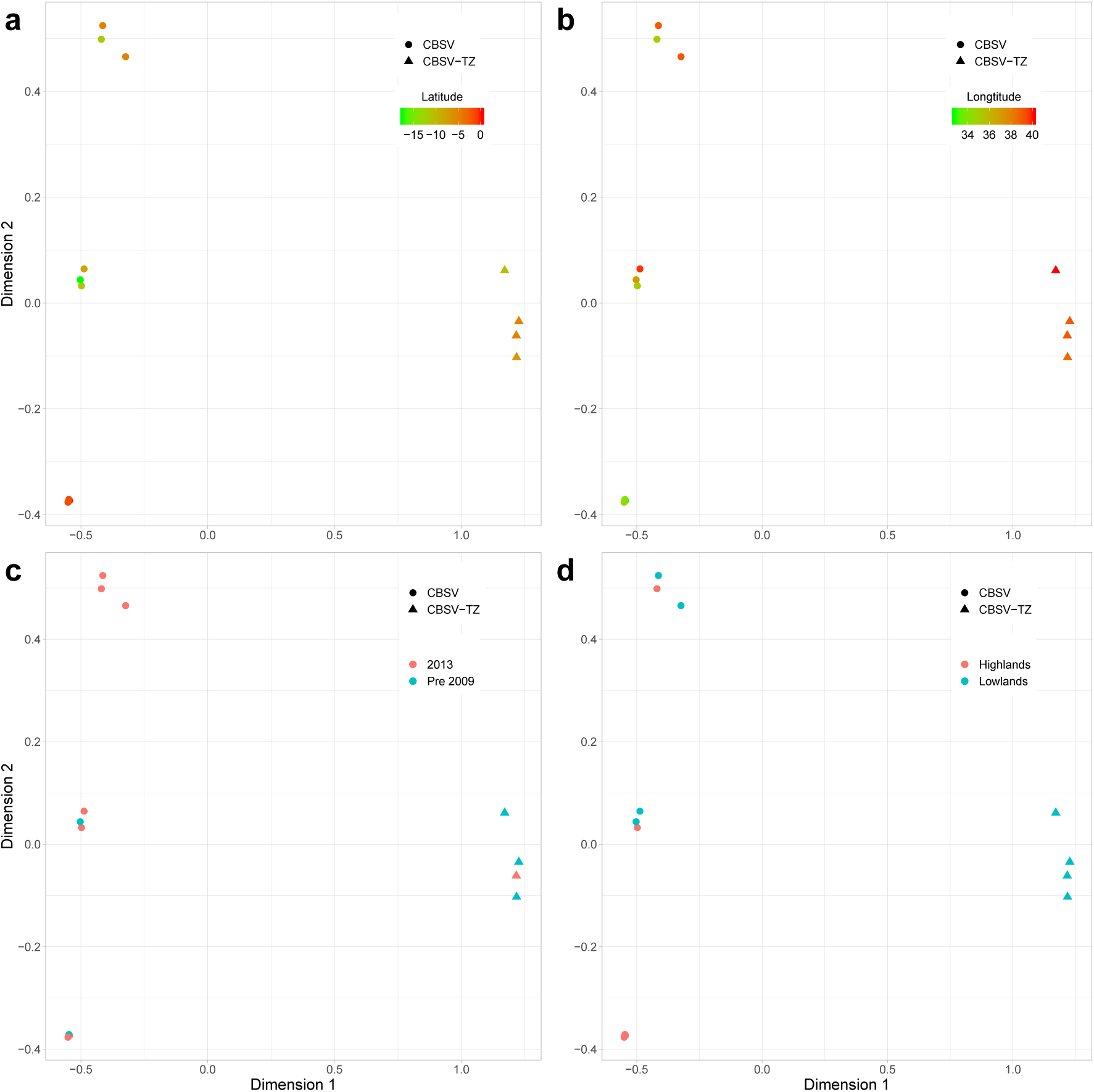
Results of MCA on the fourteen CBSV sequences. The shape of the points indicates the strain of the isolate, either CBSV or CBSV-TZ. The isolates are coloured by: a) latitude of their collection sites; b) longtitude of their collection sites; c) time of their collection (either before 2009 or 2013); and d) altitude of their collection sites (lowlands or highlands).

Figure 8 is analagous to Figure 7 except that the four CBSV-TZ isolates have been removed from the analysis. It is here that we can first see some relationship between the MCA results and the geographic origin of the isolates. Figures 8 a) and b) show that the cluster of four isolates in the top left corner of the plots are at the extreme ends of the latitude (northernmost) and longtitude (westernmost) scales. These four isolates come from the areas that surround Lake Victoria, in Uganda and Northern Tanzania.

**Figure 8:**
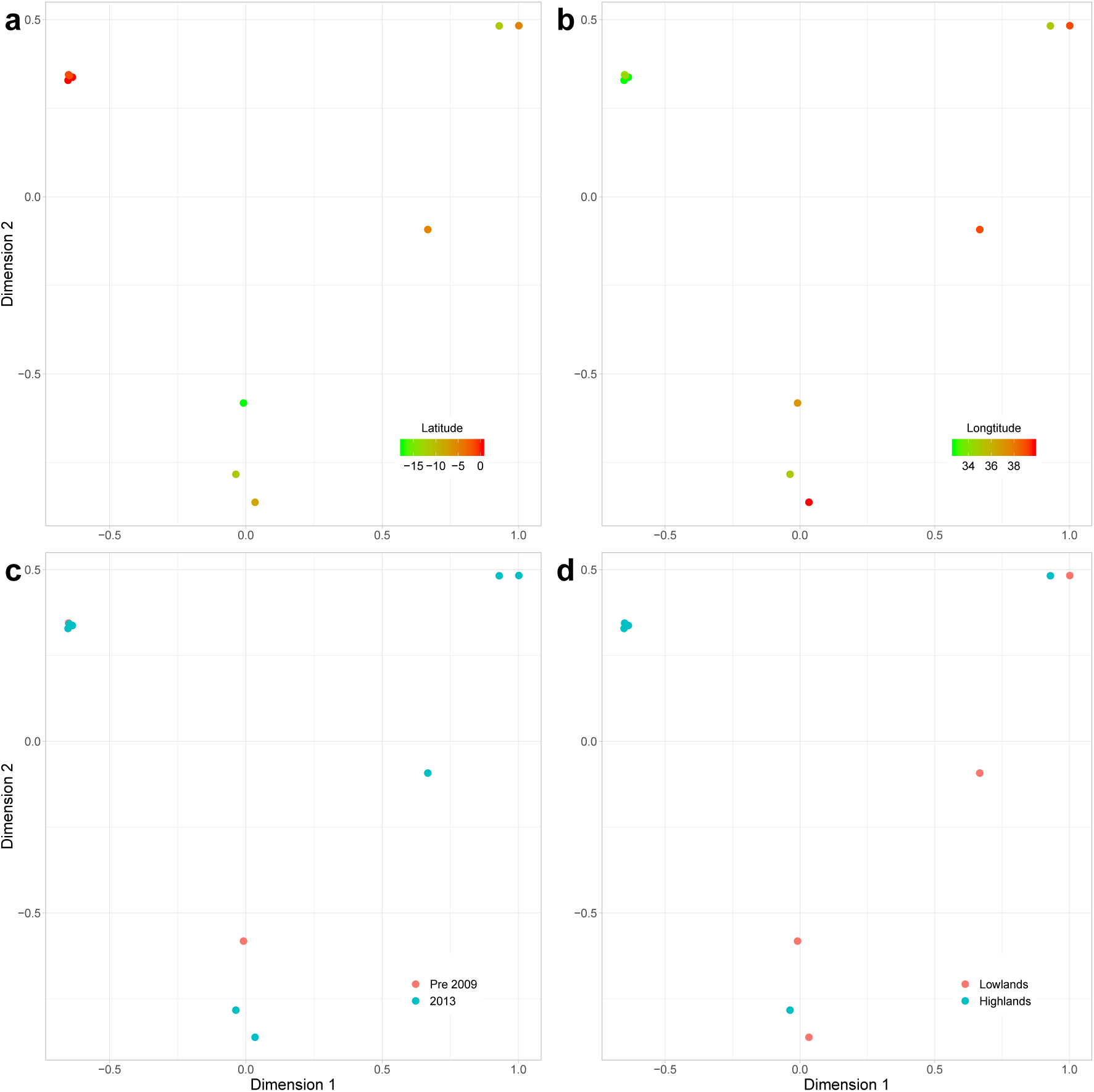
Results of MCA on the ten sequences from the CBSV strain (excluding the CBSV-TZ isolates). The isolates are coloured by: a) latitude of their collection sites; b) longtitude of their collection sites; c) time of their collection (either before 2009 or 2013); and d) altitude of their collection sites (lowlands or highlands).

Figure 9 displays the MCA results for the 15 UCBSV isolates. The most obvious feature is that two isolates strongly distinguish themselves from the remaining 13 along the first principal dimension. Even though these two isolates are clearly distinct from the others, we can see in Figure 6 that they obviously still belong to the UCBSV cluster. What is most intriguing about these two outliers, is that they appear to share very little in common. Figure 9 indicates that they are separated temporally, geographically and altitudinally. There doesn’t appear to be any other strong patterns emerging from the MCA on the UCBSV isolates. This may hint to the hypothesis that UCBSV is a more versatile virus. Isolates that are genetically very similar are able to infect plants in relatively different environments.

**Figure 9:**
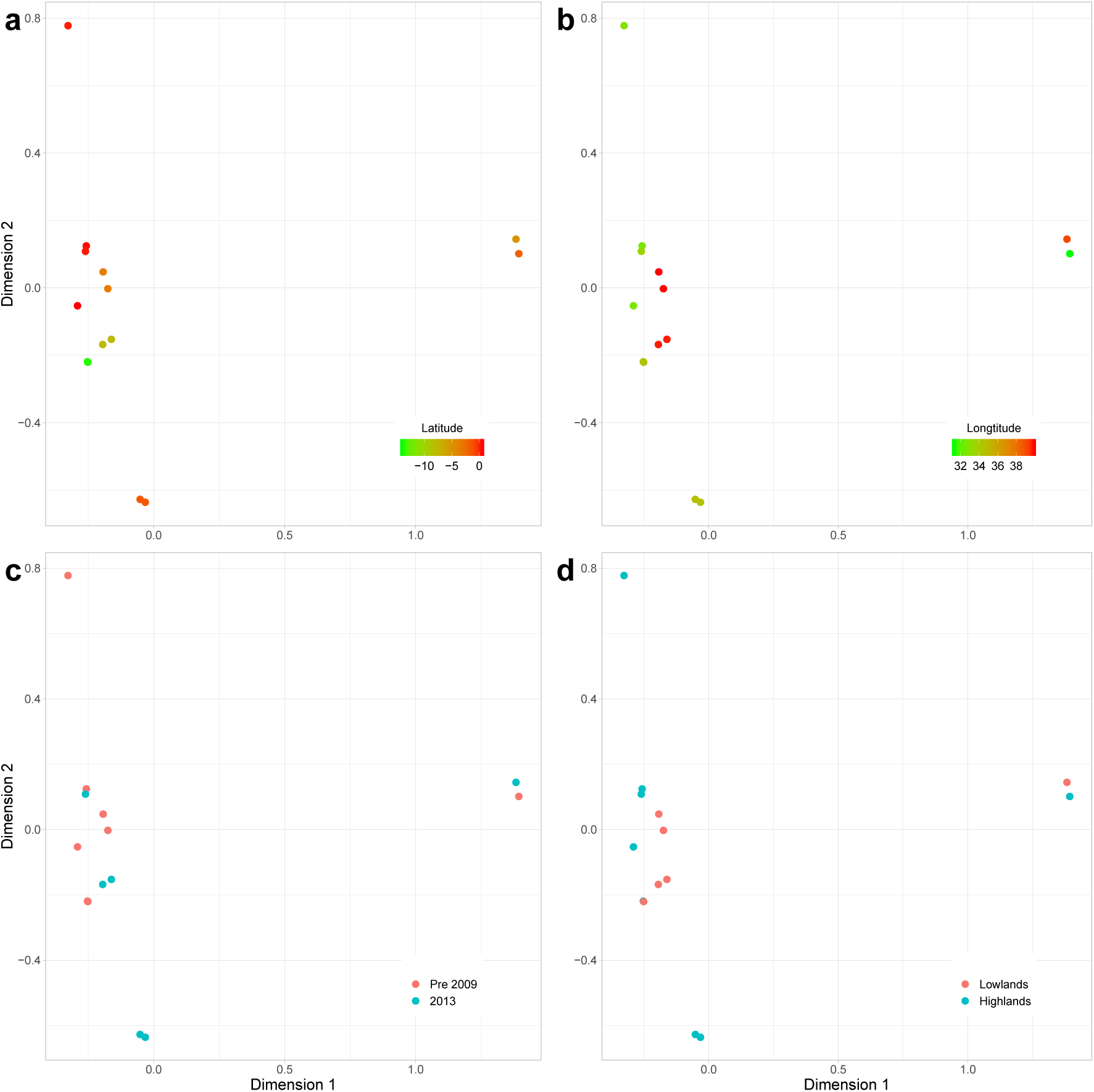
Results of MCA on the fifteen sequences from the UCBSV strain. The isolates are coloured by: a) latitude of their collection sites; b) longtitude of their collection sites; c) time of their collection (either before 2009 or 2013); and d) altitude of their collection sites (lowlands or highlands).

It must be noted here that the patterns arising from the various MCA analyses are somewhat speculative. MCA has been shown to be a powerful tool in population genetics, able to map the genetic diversity amongst a dataset to a geographical path of evolution (Rohrlach et al., 2018). However, the sparsity of our dataset ensures that strong conclusions are not possible in this case. We have 29 isolates to represent three distinct strains of a fast evolving virus, spread over a wide and diverse geographical area and collected at various intervals over a 16 year time period. Even with such a sparse dataset, we were able to notice some associations between variables that can be used to generate testable hypotheses.

## Discussion

Our results confirm and extend the work of Mbewe et al. (2017), who found evidence for a third distinct clade of CBSV by analysing a subsection of the P1 gene. Using two independent methods of sequence analysis we show that the evidence for this third clade is not restricted to the P1 gene, but is in fact present across the entire genome. That said, of the ten genes in the genome, the P1 gene is the strongest relative contributor to the signal and focusing on this gene seems apt.

There are practical implications of our findings that can have an immediate impact for cassava farmers in East Africa. The most vital step in securing the crops of farmers is the swift diagnosis of infected plants. It is vital not only to know that a plant is infected, but also to know which strain of the CBSV it has. This information is important on several fronts, it can be used to:

- select the most appropriate viral resistant cultivars for the next growing season.
- generate CBSV distribution maps that can be used to guide Agricultural Officers on where to screen (hotspot areas) or where to multiply (low pressure areas).
- inform where to deploy cassava material.
- strengthen phytosanitary regulations on movement of cassava germplasm.

This work is undertaken by scientists working in the field directly with farmers. Current practice when sequencing viral isolates is to focus on the CP gene, which can effectively distinguish between CBSV and UCBSV (Monger et al., 2001; Mbanzibwa et al., 2011). However, as indicated in Table 1, the CP gene is the least influential of the ten genes in the CBSV-TZ Class. In order to effectively and efficiently distinguish between CBSV, UCBSV and CBSV-TZ we recommend biologists focus their attention on the P1 gene, in particular the first 60 amino acids, as shown in Figures 3-5.

The results of the bioinformatic analyses presented here, when viewed in the context of the history of CBSD in East Africa, allow a speculative hypothesis of the evolution of the virus to be constructed. We know that historically, the prevalence of CBSD was negatively correlated with the altitude in which the crops were grown. The disease was prevalent in low-lying, coastal areas, and was never seen at altitudes greater than 1000m. We know from the results of the MCA that the CBSV-TZ strain (at least in the current dataset) has only been found at low altitudes, and was much more prevalent among the pre 2009 samples in our dataset than it was among the 2013 samples. Further, we know from previous research (using the same dataset as we analyse here) by Alicai et al. (2016) that CBSV is faster evolving than UCBSV. Finally, we showed that a relatively non-conserved section of the P1 gene in CBSV-TZ is strongly conserved within both CBSV and UCBSV, but strongly divergent between all three clades.

Let us denote the most recent common ancestor of the three strains as CBSV-Anc. We hypothesise that given it’s tendency to be found only at low altitudes and its apparent decreased incidence in recent years, CBSV-TZ is the closest of the three strains to CBSV-Anc. CBSV-Anc had low fitness at high altitudes, and so mutations that increased the high-altitude fitness of the virus would have been positively selected for on the fringes of its territory. Once these alleles that increase high-altitude fitness gain a foothold in the population, the geographical disparity in selective pressure may have precipitated a selective sweep of these alleles at high altitudes, but not at low altitudes. This effectively becomes the speciation event that spawns UCBSV.

As time passes UCBSV spreads through high altitude cassava growing regions where it has no competition, but also populates low altitude areas, along with CBSV-Anc. After some time the process then repeats itself. CBSV-Anc again speciates into CBSV and CBSV-TZ. Like UCBSV before it, CBSV evolved the ability to radiate into high-altitude regions. It also remains prevalent in the low-altitude regions, resulting in the reduced prevalence of CBSV-TZ, as it now competes with UCBSV and CBSV.

It can not be stated strongly enough that the above scenario is highly speculative, but it is consistent with the known history of these viruses and the evidence we can extract from the 29 whole-genome sequences. What we have highlighted however, is that there are sophisticated bioinformatics tools now available that can greatly assist in the fight against these viruses. What is needed to turn our speculations into strong and robust evidence is more data. To combat these fast-evolving viruses, we must increase research into them on two fronts:

1. We must amass much more systematic sequence data, covering the full geographic range of these viruses, with repeated sampling in each growing season. This will allow us to see in near real time how and where the virus is evolving, both geographically and on the molecular level. When sequencing virus isolates, detailed categorical information should be recorded, including the date; latitude, longtitude, and altitude of the collection site; cultivar the virus was found in; and detailed symptom description.
2. The structure of the viruses must be analysed on the molecular level. Here we highlight a region of the P1 gene that appears to be highly influential in increasing the high-altitude fitness of the virus. Understanding how and why this works on a molecular level is critical information that could assist in breeding highly resistant cultivars.

## CONCLUSION

The fight to increase food security for the world’s poorest citizens is of paramount importance. Here we show that, given a sufficient amount of data, novel bioinformatics tools have a vital role to play. Excellent work is already being done to help farmers identify and contain virus outbreaks, but these are reactive strategies. In order to eradicate these viruses we must understand how they work and evolve. For this we need much more investment in data collection and analysis. Ultimately, the cost of this investment is dwarfed by the potential gain. The benefits of achieving food security is not limited to nutrition. The resultant economic boost would allow families and countries alike to invest heavily in health and education, providing a dramatic increase to the standard of living across East Africa.

## SUPPLEMENTARY MATERIAL

**Figure S1:**
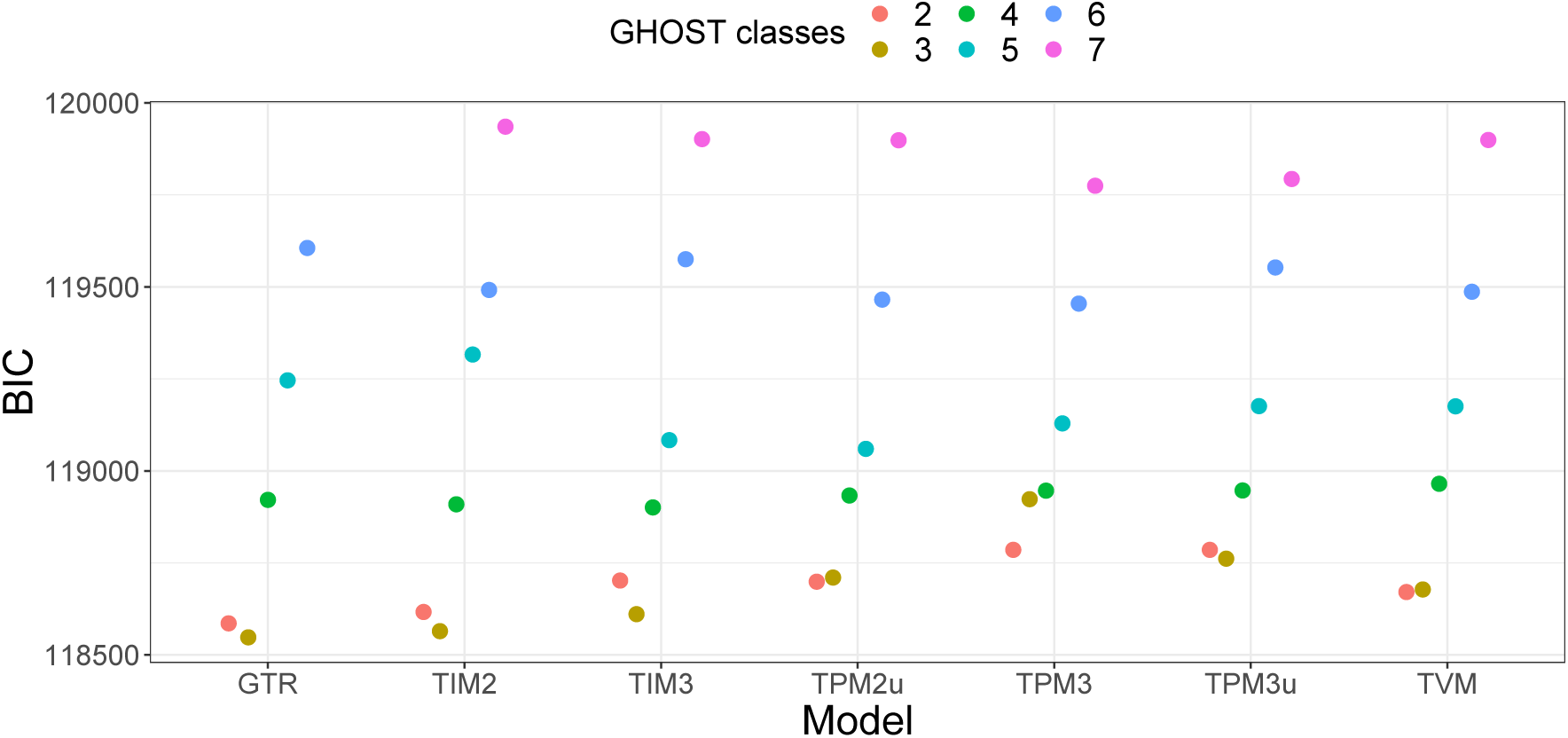
Results of model test procedure to select the optimal substitution model and number of classes, using BIC as the discriminating criterion. A total of 13 different nucleotide substitution models were tested, and each model was tested with between 2 and 12 classes. Poorly performing models (BIC > 120000) have been filtered out to improve resolution amongst the remaining models.

**Figure S2:**
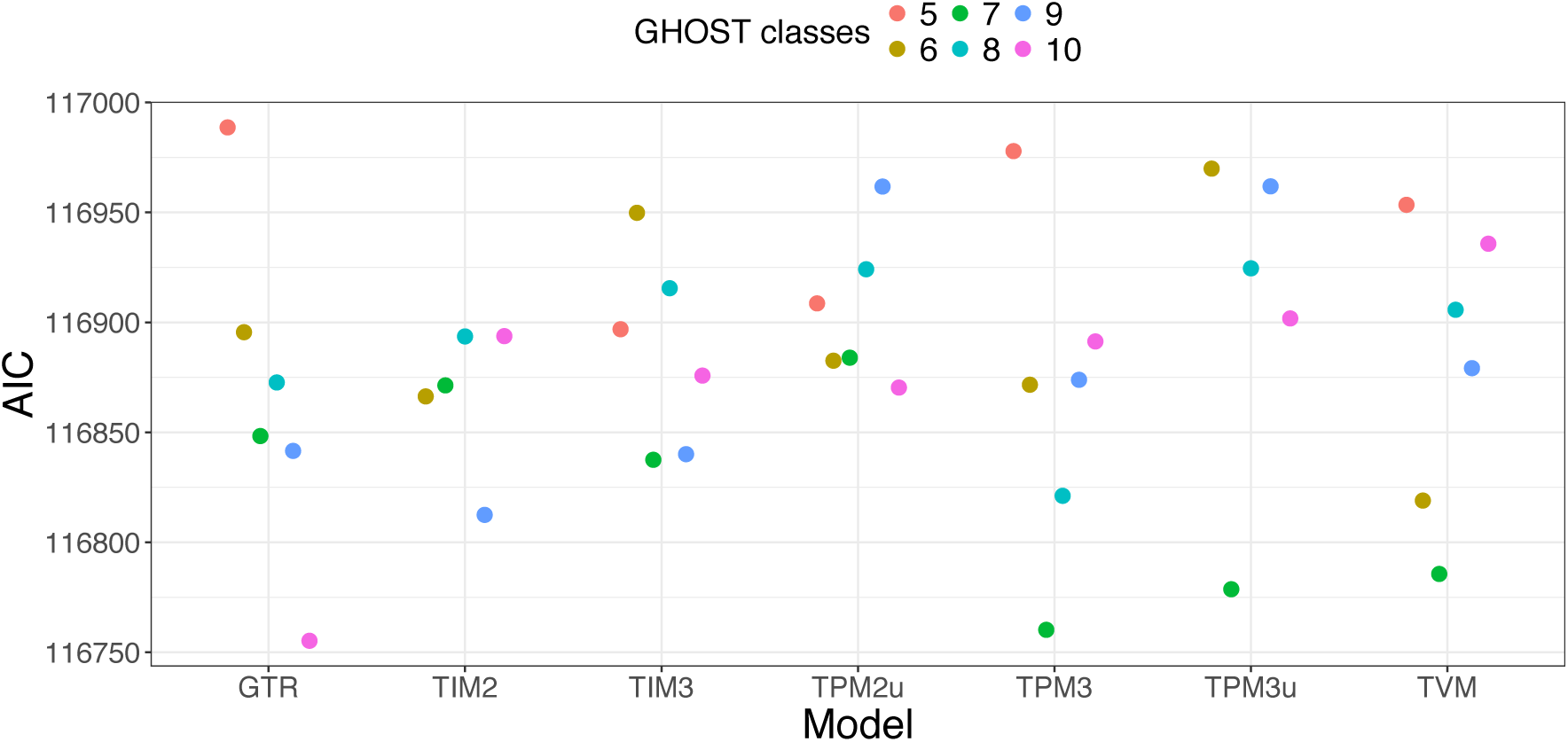
Results of model test procedure to select the optimal substitution model and number of classes, using AIC as the discriminating criterion. A total of 13 different nucleotide substitution models were tested, and each model was tested with between 2 and 12 classes. Poorly performing models (AIC > 117000) have been filtered out to improve resolution amongst the remaining models.

**Figure S3:**
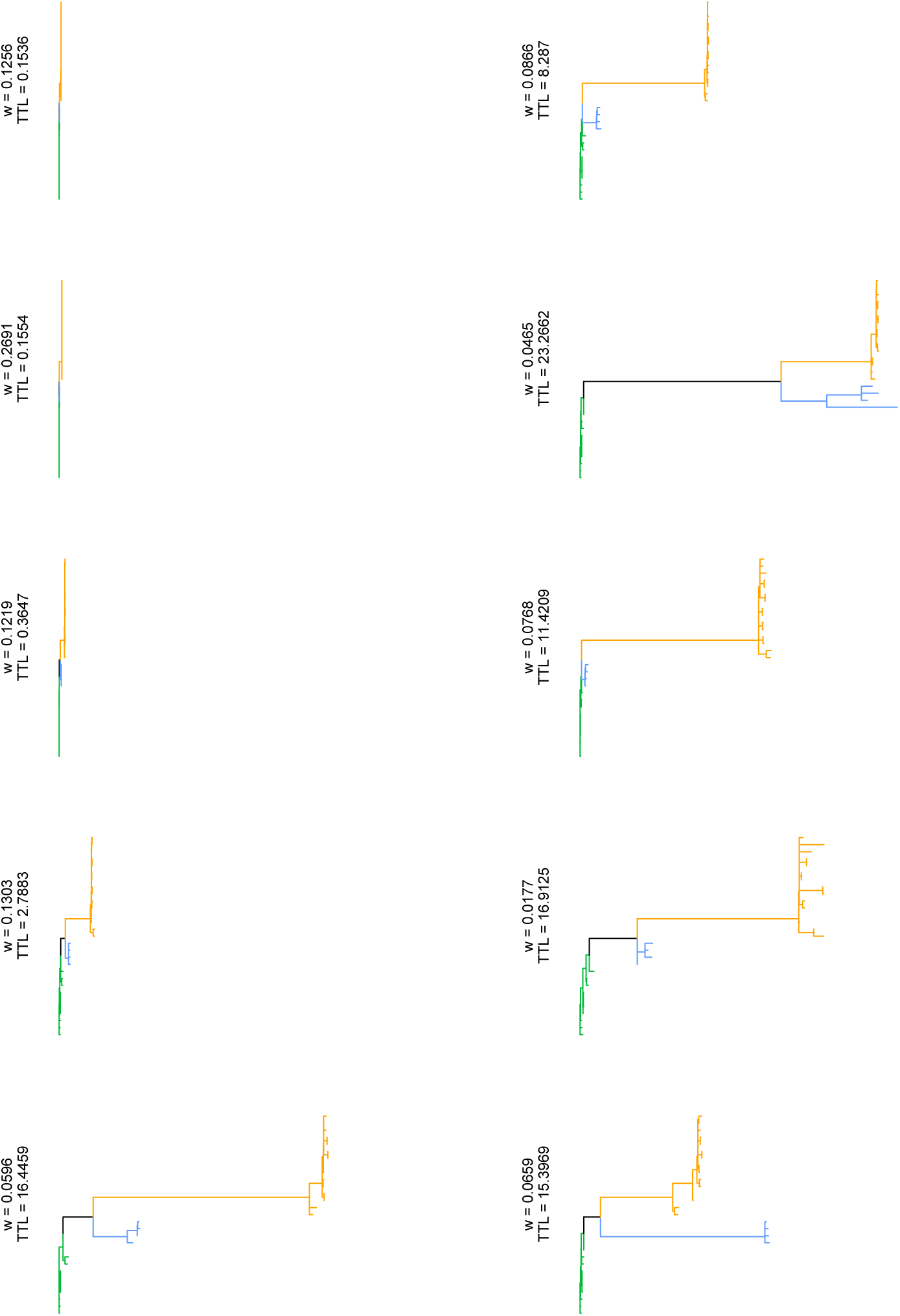
Trees inferred by IQ-TREE using the GHOST model with ten GTR classes. Tip labels are not shown for clarity. As in Figure 1, green edges form the CBSV clade, blue edges form the CBSV-TZ clade and orange edges form the UCBSV clade.

